# In-situ quality monitoring during embedded bioprinting using integrated microscopy and classical computer vision

**DOI:** 10.1101/2022.09.10.507420

**Authors:** Vasileios Sergis, Daniel Kelly, Graham Britchfield, Ankita Pramanick, Karl Mason, Andrew Daly

## Abstract

Despite significant advances in bioprinting technology, current hardware platforms lack the capability for process monitoring and quality control. This limitation hampers the translation of the technology into industrial GMP-compliant manufacturing settings. To address this, we developed a novel bioprinting platform integrating a high-resolution camera for in-situ monitoring of extrusion outcomes during embedded bioprinting. Leveraging classical computer vision and image analysis techniques, we then created a custom software module for assessing print quality. This module enables quantitative comparison of printer outputs to input CAD models, measuring area and positional accuracy. To showcase the platform’s capabilities, we then investigated how the rheological properties of granular support hydrogels impact print quality during embedded bioprinting. Our results demonstrated that lower viscosity, faster thixotropy recovery, and smaller particle sizes significantly enhance print fidelity. This novel bioprinting platform, equipped with integrated process monitoring, holds great potential for establishing robust, reliable, and auditable biofabrication processes for industrial applications.

## Introduction

Bioprinted organs promise to revolutionise medicine by tackling the organ transplant shortage and providing new animal-free platforms for drug screening and discovery. Despite significant advances, 3D bioprinting technologies, such as extrusion bioprinting, suffer from reproducibility challenges that will hamper deployment in clinical or industrial settings ^1–3^. This is due to the multitude of complex process variables that must be optimised for specific bioink compositions and bioprinting processes ^4–7^, which makes it challenging to print anatomically accurate and viable constructs consistently. Furthermore, even when optimal conditions have been identified, bioink inconsistencies and environmental fluctuations can induce extrusion errors, leading to bioprinting failures. The reproducibility challenges with existing bioprinting technology can be attributed to its inherent open-loop nature, which lacks capabilities for monitoring extrusion outputs. This technological constraint impedes the in-process optimisation and correction of material deposition parameters, which is crucial for precision during additive manufacturing processes ^8,9^. Overcoming this limitation will be important for advancing bioprinting from research to industrial and clinical applications, where producing consistent, scalable, and high-quality outputs under Good Manufacturing Practice (GMP) compliance is fundamental.

Early work in the field has shown the potential of using camera systems to monitor extrusion bioprinting outcomes ^10–12^. Such approaches can be used for in-situ assessment of print quality on the bioprinting platform. For example, an offline vision approach has been developed with a camera attached to a bioprinter to investigate the creation of a deep learning model to detect anomalies in bioprinted grid structures to determine which anomalies were most common ^13^. Similarly, integrated process monitoring and control strategies have been developed that measure feature error in bioprinted grids ^12^. The control strategy was capable of intelligently updating the extrusion inputs in the g-code to improve accuracy for future layers ^12^. While promising, cameras have typically been located at relatively large distances or oblique angles from the extrusion nozzle ^10–12,14–16^. This limits image resolution and prevents monitoring of bioink flow, filament formation, and small features within the bioprinted construct.

In this paper, we present a novel 3D bioprinting hardware prototype with an integrated camera that tracks and monitors extrusion outcomes through a transparent glass platform. This enables high-resolution process monitoring of bioink flow and filament formation during bioprinting. Additionally, we developed classical computer vision software that can compare extrusion outputs to input CAD files for offline assessment of print accuracy in terms of construct area and position. This platform was applied to investigate how variations in granular support hydrogel rheology influence print quality during embedded bioprinting, an increasingly common extrusion bioprinting modality. This novel bioprinting platform with integrated process-monitoring functionality holds great potential for enhancing quality control in bioprinting, an important consideration for facilitating the clinical and industrial translation of the technology.

## Main

### Development of bioprinting hardware with in-situ quality-monitoring capabilities

We began by developing a unique 3D bioprinting system featuring integrated microscopy for print quality monitoring. Our engineered prototype bioprinter incorporates a glass printing platform with a camera positioned directly beneath the extrusion nozzle. This novel platform enables high-resolution, unobstructed visualisation of the nozzle output area. Aligned with the extrusion nozzle, the camera’s axis performs synchronised movements, allowing it to track the printing path and visualise the bioink extrusion process in real-time (Fig 1a i-iv, Video S1). We employed a Raspberry Pi global shutter camera that can capture images at high-resolution (pixel sizes of ∼ 6 μm) and at high speed (30 fps). To enhance image brightness, we also used a camera with an illuminated lens. To test our hardware platform we employed an embedded bioprinting modality, where the bioink is extruded into a support bath, due to the increasing popularity of this method in the bioprinting community ^17–22^. In particular, the method enables precise extrusion of low-viscosity bioinks that are otherwise challenging to process using traditional on-surface extrusion printing methods. Agarose was used as our example support bath due to its relative ease of fabrication and increasing popularity for embedded bioprinting applications ^23,24^. To overcome the relative opacity of the support hydrogel we added a red dye to our bioink to help distinguish it from the surrounding support gel (Fig 1a iv). The camera system was then capable of capturing images and videos of bioink extrusion into the support hydrogel with high resolution (Fig 1a iv, Movie S1).

**Figure 1:**
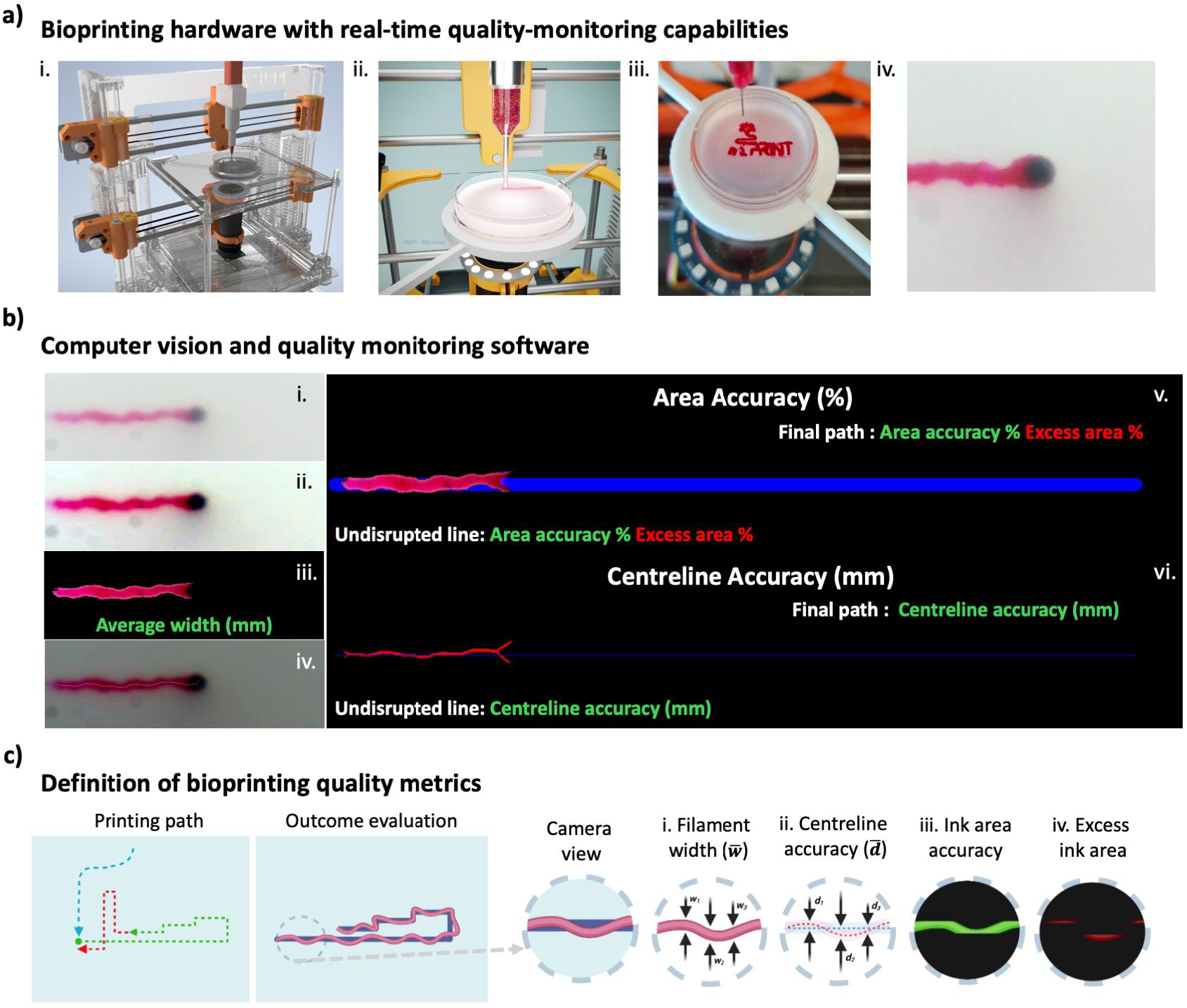
CAD model of the novel 3D bioprinter with integrated microscopy, including the bioink extruder, the elevated glass print-bed, and the bottom-view camera setup that tracks the printing process (***a***_***i***_). Illustration of the embedded bioprinter process (***a***_***ii***_) next to the physical model with petri-dish attached on a holder (***a***_***iii***_) and an example of the camera’s field of view during the 3D embedded bioprinting process (***a***_***iv***_). Illustration of the computer vision analysis steps (***b***). Image captured from the camera’s field of view with limited clarity due to formulation properties (***b***_***i***_). Enhanced image contrast, clarity, and detail acquired with percentile stretching by redistributing pixel values across the entire spectrum (***b***_***ii***_). Isolation of the printed ink using colour thresholding in HSV space, while the average width of printed structure is calculated in reference to the extracted nozzle diameter using the Hough transform technique (***b***_***iii***_). Centreline of the deposited ink extracted through skeletonization technique (***b***_***iv***_). Ink overlaid on CAD model for comparison to measure ink area accuracy and excess ink area metrics (***b***_***v***_). Ink centreline superimposed with CAD model’s centreline to measure centreline drift metric (***b***_***vi***_). The Lucas-Kanade optical flow technique was used for map creation. Top-view schematic of the print path movements and an expected printed outcome in reference to the desired CAD model (***c***). Depiction of the four metrics through an example of a zoomed-in area **(*c***_***i***_) of the expected printed outcome: average printed structure width **(*c***_***ii***_), centreline drift of the physical printed path in reference to the CAD design **(*c***_***iii***_), ink area accuracy of the deposited ink overlapping the CAD model **(*c***_***iv***_), excess ink area outside of the CAD design **(*c***_***v***_).

Camera systems have been previously employed to monitor bioprinting outcomes, but typically at oblique angles and/or larger distances from the extrusion nozzle ^10,11,14–16^. Our unique axis cloning approach positions the camera directly beneath the extrusion nozzle during the bioprinting process, enabling high-resolution imaging and tracking of bioink flow. Notably, as the camera can move in synch with the extrusion nozzle, this opens up the possibility of stitching multiple images together to reconstruct large areas at high spatial resolutions. As will be demonstrated in the next section, this innovative bioprinting hardware offers a robust platform for quantitative quality monitoring.

### Development of quality monitoring software using computer vision techniques

Next, we developed a workflow for quantifying print quality from video recordings of the extrusion process using image processing and classical computer vision methodologies. Firstly, the contrast, clarity, and detail of the extruded ink (in red) was enhanced using percentile stretching by redistributing pixel values across the entire spectrum (Fig 1b_ii_). The printed ink was then isolated from the image using colour thresholding in HSV space, and the average width of extruded bioink was calculated in reference to the extracted nozzle diameter using the Hough transform technique (Fig 1b_iii_). Next, a centreline of the printed filament was extracted through a skeletonization technique and overlaid with the original image (Fig 1b_iv_). The printed ink area was then overlaid onto the input CAD model to assess print accuracy. The percentage overlapping/excess ink areas were then assessed to quantify print accuracy (Fig 1b_v_). The platform was also used to measure the average distance between the centrelines of the ink and CAD model to assess print path accuracy (Fig 1b_vi_). Lucas-Kanade optical flow technique was used to measure the average optical flow of the ink for map creation. This novel software platform enables 2D reconstruction of bioink extrusion, which has not been reported before. Importantly, the platform also enables direct comparison of bioink extrusion outputs to the input CAD files for quality monitoring assessments.

Leveraging this software platform, we next established quantitative metrics to assess bioprinting extrusion quality. To provide a quantitative 2D assessment of print quality, we defined four metrics including the 1) average width of the printed filament, 2) centreline drift (the average distance between the printed structure’s centreline and the desired print path), 3) ink area accuracy (the percentage of overlapping area between the extruded bioink and the CAD model) and 4) excess ink area (percentage of area covered by excess extruded ink). These metrics enable quantitative evaluation of the bioprinting process outcomes, specifically in terms of area and positional accuracy of the printed structure. This form of quality monitoring has not been previously reported. The quality metric analysis is processed offline using videos of the extrusion process, and the computational time for the image analysis process is approximately 0.5 seconds per frame. Through future iterations, it may be possible to use this platform for real-time evaluation and display of quality metrics to the user during the bioprinting process. This platform holds great potential for diverse quality monitoring applications in bioprinting.

In terms of current limitations, it should be noted that the platform quantifies print outcomes based on simplified 2D representations of the bioprinting process. This may be sufficient for many quality monitoring applications, but future work will explore integrating additional camera vantage points for 3D reconstruction of bioink extrusion. Additionally, our system relies on the colour of the bioink being distinct from the surrounding support bath, although a variety of dyes could be employed. The transparency of the support bath will also influence the ease and accuracy of the bioink reconstruction process. For the agarose support bath formulations used in this study, we were able to image the bioink at depths of up to 3 mm. Finally, as mentioned above, the analysis is processed offline on video recordings of the extrusion process, but future work will explore real-time versions of the method. While this work focused on embedded bioprinting, the system can also be applied to traditional on-surface 3D bioprinting (without a support bath).

### Rheological characterisation of granular support hydrogels with varied particle morphology

As mentioned previously, embedded bioprinting, a technique where the bioink is extruded into a secondary support hydrogel, is an increasingly common bioprinting modality. Support hydrogels, typically engineered to exhibit Bingham plastic behaviour, act as a solid at low shear stress but flow like a liquid under higher stress. This property allows local transformation from a solid-like state to a liquid-like state around the nozzle during extrusion, followed by transformation back to a solid-like state after the bioink deposition, thereby enclosing the deposited bioink to maintain structural shape and position ^25^. Quality outcomes during embedded bioprinting are therefore dictated by the rheological properties of the support bath. While previous research has explored the link between support bath rheological properties and print quality ^16,18,26^, consistent and generalisable relationships have yet to be established. To address this gap in knowledge, we leveraged our novel hardware platform to investigate how the rheology of granular support hydrogels influences key print quality metrics during embedded bioprinting.

First, we set about establishing a range of support bath formulations with varied rheological properties. Our agarose-based supports are fabricated using a shearing method that transforms a cooling agarose solution into a granular material with shear-thinning and self-healing properties ^27,28^, so we therefore tested how varying polymer concentration and shearing rate influenced structure and rheological behaviour. Interestingly, increasing the stirring speed from 350-1400rpm resulted in distinct differences in particle size and morphology (Fig 2a i). The particles broadly shared a similar overall structure - a central mass making up most of the body, with elongated protrusions extending from this body. Notably, we observed pronounced differences in the size of the particle body, with significantly smaller particle sizes for higher polymer concentrations and stirring rates (Fig 2a ii). Particle size is an important consideration for granular support baths as it will strongly influence the resolution and smoothness of the bioprinted structures. The particles also displayed distinct protrusions from the central body that likely influence rheological properties through physical particle-particle interactions. These distinct particle morphologies have been reported before, and are thought to be caused by temperature gradients that arise as shear is applied to the material as it crosslinks during cooling ^28–30^.

**Figure 2:**
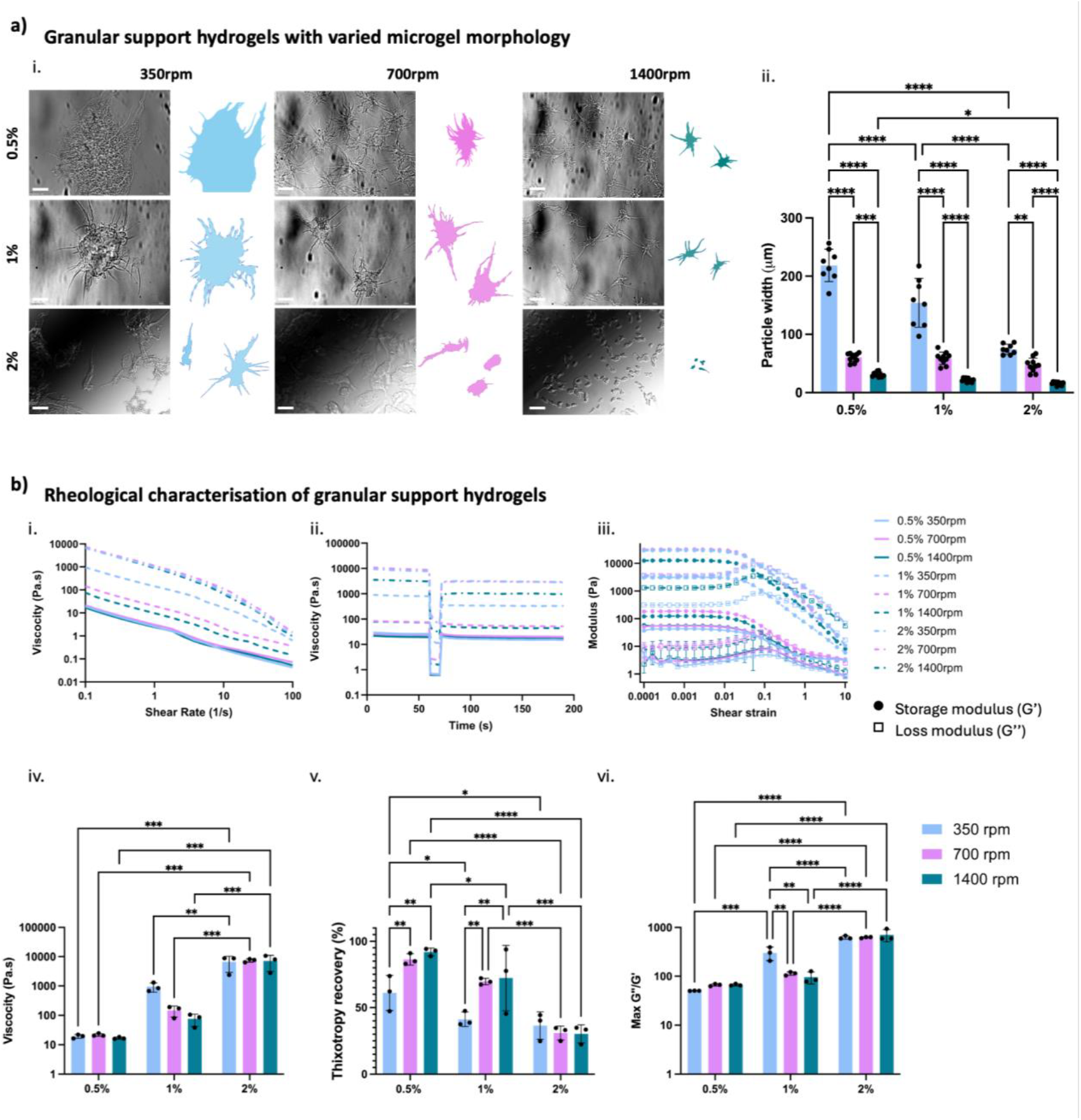
Morphological and rheological characterisation of the support bath hydrogel formulations. Images captured using a brightfield microscope of the various support hydrogel formulations (scale bar 50*μm*). Schematics of the particles extracted to facilitate the comparison of particle morphology within the support hydrogels formulations (***a***_***i***_). Measurements are for average particle width. Bar graphs show n = 8-10 individual particles, mean ± s.d, two-way ANOVA (n.s. not significant, *p < 0.05, **p < 0.01, *** p < 0.001, **** p < 0.0001) (***a***_***ii***_). Shear rate sweeps test with all solutions demonstrating shear-thinning behaviour (***b***_***i***_), and relative bar graph comparing the viscosity of the gels in low strain (***b***_***iv***_). Three interval thixotropy test to assess the structural recovery of the formulations(***b***_***ii***_), while the relative bar graph summarizes the recovery in percentage (***b***_***v***_). Finaly, oscillatory amplitude sweep tests performed to investigate the elastic and viscous properties (***b***_***iii***_), with the bar graph presenting the loss to storage ratio of the formulations (***b***_***vi***_). Bar graphs show n = 3 technical replicates, mean ± s.d, two way ANOVA (n.s. not significant, *p < 0.05, **p < 0.01, *** p < 0.001, **** p < 0.0001).

In terms of rheological properties, shear rate sweeps demonstrated that all formulations displayed shear-thinning properties (Fig 2b i). As expected, viscosity increased for higher polymer concentrations (Fig 2b i, iv). Interestingly, higher stirring rates trended towards reduced viscosity at medium polymer concentrations, potentially due to changes in particle morphology (Fig 2b i, iv). Thixotropy tests demonstrated that all support hydrogels displayed self-healing properties as they were cycled through periods of low-high-low strain (Fig 2 b ii). We quantified the extent of self-healing by measuring the percentage viscosity recovery through the strain cycles, with greater recovery observed for the formulations produced with higher stirring speeds at low and medium polymer concentrations (Fig 2 v). Lower levels of thixotropic recovery were observed for supports produced with higher polymer concentrations (Fig 2 v). Oscillatory amplitude sweeps demonstrated that storage and loss modulus values increased for higher polymer concentrations (Fig 2b iii). Yielding to more fluid-like behaviour was also observed at higher shear strains, further confirming shear-thinning properties. Formulations with lower polymer concentrations did not exhibit a clear yield point, defined as the crossover between the storage and loss modulus. Due to the absence of a yield point for some formulations, the ratio of loss to storage modulus was compared between the groups (Fig 2b vi). Interestingly, an increase in this ratio was observed as the polymer concentration increased, potentially attributed to the greater resistance to deformation at higher polymer concentrations or the smaller particle size (Fig 2b vi). After comprehensive rheological characterisation of the various support bath formulations, we proceeded to investigate the impact of these properties on print fidelity and resolution during embedded bioprinting.

### In-situ quality monitoring during embedded bioprinting in granular support hydrogels

While particle-based support hydrogels are increasingly utilised in embedded bioprinting, a comprehensive understanding of how their rheological properties affect print outcomes has not been established. Previous research indicates that higher support hydrogel viscosity can improve print resolution, measured by quantifying the width of extruded filaments ^16,18,26^. Conversely, lower viscosity can cause greater filament instability and buckling during deposition, particularly when the extrusion and nozzle translation speeds are not matched ^16^. To help better understand the impact of rheological properties on print quality, we used our novel bioprinting hardware with integrated quality monitoring to assess how variations in support bath rheology influenced embedded bioprinting outcomes. In particular, we were interested in investigating how the fluidic behaviour of the support hydrogel could impact more complex aspects of the printing process, such as direction changes and the disruption of previously deposited ink during subsequent deposition steps. To do this, we designed a print path to assess how successive printing steps, including parallel and perpendicular motion at increasing proximities, could influence our quality metrics (Fig 1c). Disruption of previously bioprinted structures can occur during successive printing layers, the extent of which is likely influenced by the rheological response of the support hydrogel. Shear forces from the moving nozzle and hydrodynamic forces from fluid flow can displace the hydrogel particles from their original positions, with subsequent repositioning facilitated to varying degrees depending on the thixotropic behaviour of the material. This disruption may occur due to inner tension within the hydrogel, which arises from the mechanical stresses and forces exerted during the printing process.

Subsequently, we conducted print quality tests for all support hydrogel formulations, utilizing our quality-monitoring platform to quantitatively compare outcomes (Fig. 3). The platform first converted brightfield images into comprehensive print quality maps, detailing the centreline and area of the printed ink alongside a direct comparison to the input CAD file (Fig. 3). Print outcomes captured by the camera were stitched together to create a single high-resolution map of the entire print path (Fig. 3a). Centreline accuracy maps were generated by mapping the discrepancy between the desired CAD model path (blue line) and the actual printed path (green line) (Fig. 3b). We also differentiated the actual print path centreline both before and after bioprinting an additional filament adjacent to the initial filament, assigning distinct colours to the print paths after each stage to visualise shifts in location (Fig. 3b). The centreline was assigned red for the initial filament and then switched to green after the second filament was printed. This resulted in regions of yellow appearing when these two lines overlapped. The software also calculated the distance between the centrelines of the CAD model and the printed structure after deposition of the first filament (undisrupted line) and the second filament (final path). The platform also displayed both ink accuracy and excess ink areas relative to the input CAD model (Fig. 3c). Bright magenta indicates areas where the initial and second filaments overlap, while faded magenta represents the full print path where the initial line is absent (Fig. 3c). This analysis allowed us to assess whether printing the second filament influenced the position of the first. The software also automatically provided percentage values for ink area accuracy (green) and excess ink area (red) (Fig. 3c). These values were quantified both after deposition of the first (undisrupted line) and second (final path) segments, revealing the impact of subsequent printing on the initial filament.

**Figure 3:**
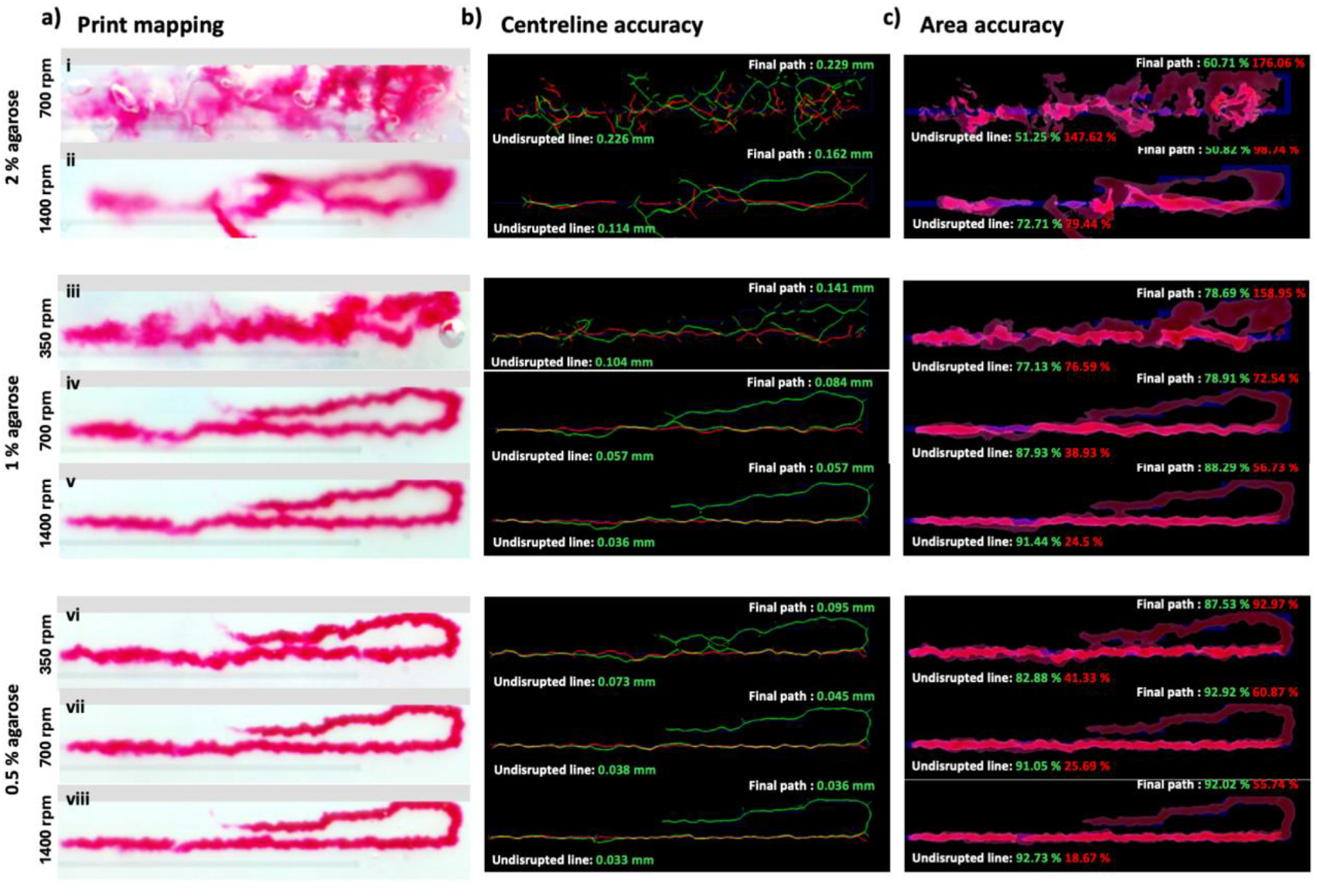
Illustration of print outcomes using agarose support baths with concentrations of 0.5%, 1%, or 2% agarose particles, prepared with stirring rates of 350 rpm, 700 rpm, or 1400 rpm. Mapped result of frames captured from the camera and stitched together (***a***). Result of the centreline drift outcome between the desired CAD model path and the actual printed path, with the blue line representing the target path’s centreline, the red line representing the initial filaments centreline (undisrupted), and the green line representing the final print path’s centreline after the second filament was printed. Note, regions of yellow appear where the red and green centrelines overlap. The software also displays the distance between the centreline of the CAD model and the printed structure after printing the first filament (undisrupted line) and the second filament (final path) (***b***). Results of the ink accuracy and excess ink areas in reference to the desired CAD model path and width (***c***). The ink area is magenta, and the CAD model area is blue. Both the first and second filaments were assigned magenta, and the brighter regions therefore represent the overlap between the first and second filaments. The faded magenta areas show the full path minus the initial filament.

Overall, we observed significant improvements in print quality metrics for lower polymer concentrations (Fig 3). Notably, consistent improvements in print quality were observed for formulations manufactured at 1400rpm, potentially attributable to their smaller particle size (Fig 3). Despite possessing shear-thinning and self-healing properties, accurate embedded bioprinting proved highly challenging for the 2% 350 and 700 rpm formulations (Fig 3), so these groups were excluded from full quantitative analysis. All other support bath formulations underwent triplicate print quality testing (Fig. 4a). Lower concentration formulations yielded the best print quality outcomes, with the 0.5% groups demonstrating the highest ink accuracy area (Fig 4a iii). Similarly, these groups exhibited the lowest levels of centreline drift and excess ink area (Fig 4a ii, iv). Additionally, lower concentration formulations produced the narrowest average filament width, converging towards the targeted nozzle width of 285 microns (Fig 4a i). The 0.5% concentration fabricated at 1400 rpm achieved the closest match to the target, deviating by only 19 microns (Fig. 4a i).

**Figure 4:**
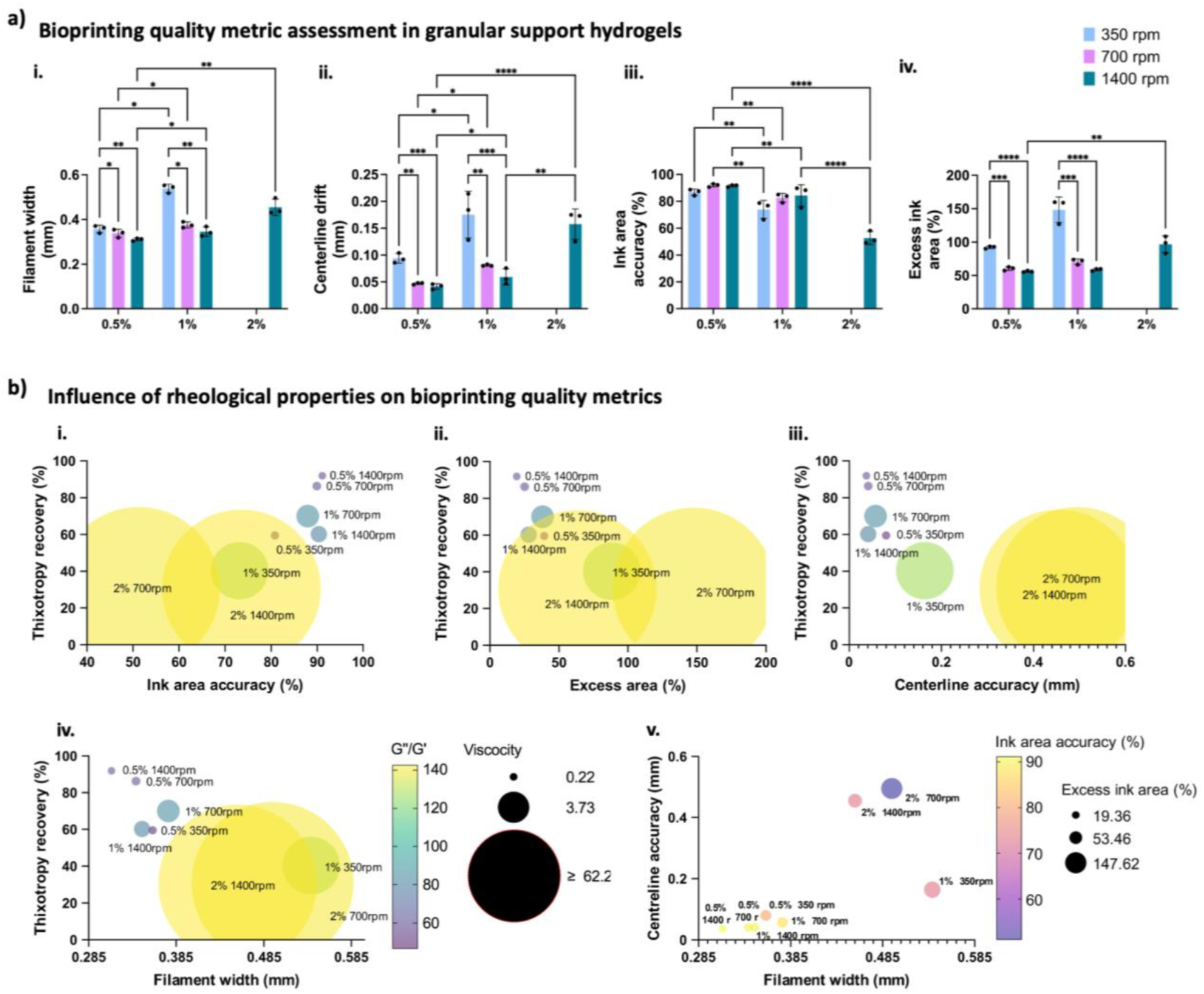
(***a***) Bar graphs summarising the results for each metric individually including filament width (***a***_***i***_), centreline drift (***a***_***ii***_), ink area accuracy (***a***_***iii***_), and excess ink area (***a***_***iv***_). Bar graphs denoted by n = 3 technical replicates, mean ± s.d, two-way ANOVA (n.s. not significant, *p < 0.05, **p < 0.01, *** p < 0.001, **** p < 0.0001). Note, bar graphs for 350rpm and 700rpm at 2% have been excluded from the analysis as it was not possible to get discernible measurements for n=3 samples. (***b***) Multivariable graphs combining print quality metrics with the rheology data of the support bath formulations. Each formulation is presented in the form of a circle, with the diameter denoting the viscosity level, the colour the loss-to-storage modulus ratio value, and the Y axis is the thixotropy recovery. Ink area accuracy (***b***_***i***_), excess ink area (***b***_***ii***_), centreline drift (***b***_***iii***_), and filament width (***b***_***iv***_) are presented on the x-axis. Finally, all metrics are presented together in a single multivariable graph with each formulation presented in the form of a circle, with the diameter denoting the excess ink area level, the colour the ink area accuracy value, while the X axis the structure’s average width and the Y axis the centreline drift (***b***_***v***_).

For a given polymer concentration, the stirring rate was also found to have a significant influence on print quality metrics. For example, for the 1% polymer concentration, higher stirring rates led to substantial improvements in print quality. This was evident in reduced centreline drifts and excess ink areas (Fig 4a ii, iv), narrower filament widths (Fig 4a i), and increased ink area accuracy (Fig 4 a iii). These findings highlight how relatively small changes to the support bath manufacturing process can significantly impact print outcomes. The observed results are likely attributable to the smaller particle sizes achieved with higher stirring rates. In general, smaller particles correlated with reduced centreline drift, approaching the CAD model’s intended path. Conversely, granular supports formed using larger particles resulted in substantial print distortions, with deformed path trajectories that deviated significantly from the desired outcome. Similar clear trends emerged concerning the excess ink area. Lower stirring rates (e.g., larger particles) resulted in inaccurate bioink deposition, deviating from the CAD model’s targeted path. The support bath formulation with the lowest percentage of excess ink area, below 19%, was the 0.5% concentration at 1400 rpm (Fig. 3b viii).

As mentioned previously, print quality metrics for embedded printing are strongly dependent on the rheological properties of the support bath. Aiming to reveal relationships between these factors, we next plotted multivariable representations of our four print quality metrics as a function of rheological properties (Fig 4). Out of the seven formulations, the 0.5% concentration formed at 1400 rpm had the best overall performance across all metrics (Fig 4b). In terms of rheological behaviour, this formulation combined lower viscosity, lower loss-to-storage modulus ratio, and higher thixotropy recovery. The multivariable representations broadly demonstrated that support bath formulations with these properties produced better print outcomes (Fig 4b i-iv). This combination of properties resulted in higher ink area accuracy, with the best solutions positioned in the top-right corner of the graph (Fig 4b i). These properties also resulted in the lowest excess ink area (Fig 4b ii) and the lowest centreline draft (Fig 4b iii), with the best solutions located on the left side of the graph. Finally, these properties also produced the closest match to the targeted filament width, with the best solutions again located on the left of the graph (Fig 4b iv). We then combined our four print quality metrics on a single multivariable representation for comparison between the different formulations (Fig 4b v). With the aim of 1) reducing the centreline drift, 2) increasing the ink overlap area, 3) reducing the excess ink area, and 4) matching the designed CAD model filament width, the best support bath solutions are positioned at the bottom left corner of the graph (Fig 4b v).

Our results provide a comprehensive analysis of how the rheological properties of support baths influence print quality metrics in embedded bioprinting. Notably, we found that lower viscosity formulations led to improved print accuracy. This finding is somewhat in contrast to previous literature reporting increased print accuracy with higher viscosity agarose support hydrogels ^16^. These contrasting findings may be due to differences in the viscosity ranges investigated and variations in support bath preparation protocols. The observed improvement with lower viscosity formulations could be due to reduced peripheral material flow during nozzle movement, which may be more likely to occur with higher viscosity granular hydrogels and disrupt previously printed structures. Considering the particle morphology analysis, our results also suggest that print quality improvements may be linked to changes in the constituent particles’ size and shape. Notably, the best-performing support hydrogel formulation had relatively small particle sizes (without being the smallest), featuring multiple protrusions connected to the central body of the particle. Previous literature has reported improved print quality outcomes for support hydrogel formulations with smaller particle sizes ^18,19,31^. Additionally, our findings indicate that support hydrogel formulations with enhanced thixotropy recovery (or self-healing properties) can improve print accuracy. This result is expected, as the self-healing properties of the support hydrogel allow it to reform around the nozzle during printing, aiding in the precise localisation of the bioink in 3D space. Altogether, our results indicate that within the range of agarose support bath formulations tested, those with lower viscosity, faster thixotropy recovery, and smaller particle sizes yield superior print accuracy in embedded bioprinting. These findings contribute to a more generalised understanding of the relationship between rheological properties and print accuracy in embedded bioprinting and can be leveraged to inform the design of future support bath formulations.

## Conclusion

We successfully engineered a novel bioprinting platform that enables high-resolution in-situ process monitoring of extrusion outcomes. The platform includes innovative software that enables precise comparison of printed outputs to input CAD models for comprehensive quality assessment. We showcased the platform’s ability to assess key print quality metrics (area and positional accuracy) during bioprinting process development, providing users with advanced quality assessment data that can be used to optimise workflows. Such functionalities will be essential for translating bioprinting technology into GMP-compliant environments.

## Supporting information

Video S1 - Print quality analysis for 0.5% agarose 1400rpm formulation

## Acknowledgements

This publication has emanated from research conducted with the financial support of the EU Commission Recovery and Resilience Facility under the Science Foundation Ireland Future Digital Challenge Grant Number 22/NCF/FD/10991. This publication has also emanated from research supported in part by a grant from Science Foundation Ireland and is co-funded under the European Regional Development Fund under Grant number 13/RC/2073_P2.

## Author Contributions

Conceptualisation and design, V.S., D.K., K.M., and A.D.; Execution of experiments V.S., D.K., G.B., A.P.; Writing, review, and editing, V.S., D.K., G.B., A.P., K.M., and A.D.; Supervision, A.D.;.

## Methodology

### Bioink and support hydrogel formulations

In this study, agarose particles were chosen as a proof of concept among commonly reported materials for support hydrogel particle formulations to test the system and methodology. The agarose solutions were prepared using the shearing method, a widely used method for creating granular support hydrogels ^24,26,32^. This method entailed dissolving agarose powder (Type I, low EEO, Sigma Aldrich) in PBS at concentrations of 0.5%, 1%, or 2%. Subsequently, each solution underwent sterilisation via autoclaving (Prestige Medical Classic Autoclave, Fisher Scientific), to ensure both sterility and uniform melting of the agarose to achieve a homogenous solution. The melted agarose solution was then subjected to varying speeds on a magnetic stirrer, including 350 rpm, 700 rpm, and 1400 rpm, for 3 hours while gradually cooling to ambient room temperature. This process fosters the formation of a suspension of agarose particles as the polymer cools under shearing ^28–30^. The variation in stirring rate and agarose concentration resulted in nine different formulations following a three-level two-factor full factorial design. The agarose solutions were then added to a glass bottom petri dish for embedded bioprinting, as seen in Fig. 1, allowing in-line monitoring of the printing process. Regarding the bioink used, alginate was selected due to its widespread use for bioprinting applications. The ink was prepared by dissolving alginic acid powder (alginic acid sodium salt from brown algae, Sigma Aldrich) in phosphate-buffered solution (PBS) at a concentration of 7wt%. To enhance contrast and visualisation of the printed structure against the background formed by the agarose support bath formulations, a red dye (Direct Red 80, Sigma Aldrich) was added at a concentration of 0.1 mg/ml. Fig. 1a iv depicts an example of the camera field of view during the printing process, with clarity variation of the deposited ink depending on the support bath solution.

### Rheological characterisation

Rheological characterisation was conducted to assess the viscoelastic properties of the hydrogel support bath solutions using shear rheometry (TA Instruments AR2000, cone and plate geometry). This comprehensive analysis included amplitude sweep, viscosity curve, and thixotropic analyses to evaluate key rheological properties relevant to the embedded bioprinting process. The testing encompassed seven formulations with three repetitions performed on each. Viscosity sweeps were conducted within a shear stress range of 0.1-100 1/s, while a shear strain sweep ranging from 0.0001-10 was utilised to characterise storage modulus (G’) and loss modulus (G’’). To compare the solutions effectively, the loss to storage modulus ratio was calculated, along with the assessment of the descending slope of the storage modulus and the point where it initiated descending after the linear viscoelastic (LVE) region, indicative of the structural decomposition due to increased shear stress. Furthermore, a three-interval thixotropic analysis was employed to assess structural regeneration or self-healing capabilities of each solution. This involved the application of low shear for 60 seconds (0.01), followed by high shear for 10 seconds (0.1), and concluding with low shear for 120 seconds in the third interval (0.01). Viscosity measurements were recorded throughout each strain cycle to capture the material’s rheological response under varying conditions.

### 3D Bioprinter with integrated microscopy

The bioprinter developed in this study introduces a novel approach for in-line monitoring of the freshly extruded bioink layer within an embedded bath, utilising a bottom-view camera setup. The developed bioprinter integrates a set of electronic components, including microprocessors, actuators, and a camera, while leveraging the spatial movement of a pre-existing desktop printer. The image data collected from the mechanism is processed offline, taking approximately 0.5 seconds to generate the detailed print quality reports using the classical computer vision and image processing methodologies. The CAD model of the developed bioprinter is presented in Fig. 1a. The overall dimensions of the printer are approximately 500×550×400 mm in length, width, and height. Notably, several modifications have been made to the printer, one of which involved replacing the existing extruder with a progressive cavity pump (Puredyne™, Germany). This modification was aimed at enhancing control over the flow rate of the ink, crucial for precise bioprinting applications. When compared to alternative bioprinting extrusion methods such as pneumatic, piston-like, or auger screw extruders, the progressive cavity pump can maintain precise control over ink deposition, regardless of the ink’s rheological properties. Additionally, the precisely regulated volumetric displacement of the progressive cavity pump helps minimise shear forces exerted on bioinks, thereby enhancing cell viability. To address concerns about visibility and potential obstructions in monitoring the printing process, the decision to position the camera underneath a glass bed table was made. This configuration ensures an unobstructed view of the entire surface, optimising visibility for reliable monitoring. To mount the camera, a space beneath the table was necessary. Consequently, a platform comprising four threaded studs and custom 3D-printed components was incorporated to support a thin square-shaped glass. The camera is affixed to an axis parallel to the printer’s main extrusion axis, ensuring aligned movement during printing. This synchronised motion allows the camera to closely monitor the bottom view of the freshly extruded ink, as depicted in Fig. 1. By moving in tandem with the extruder, the camera captures accurate real-time data of the printing process, enhancing monitoring precision. The Raspberry Pi Global Shutter Camera was chosen for its capability to capture rapid motion without the artefacts typically associated with rolling shutter cameras. Featuring a 1.6-megapixel Sony IMX296 sensor with a pixel size of 3.45μm × 3.45μm, it excels in fast-motion high-speed photography, boasting high light sensitivity and requiring short exposure times. To enhance the camera’s imaging capabilities, a 100X industrial microscope lens has been integrated. This lens provides magnification for detailed and accurate imaging, offering a C/CS-mount for seamless integration. Additionally, a 16xRGB LED Ring Board has been integrated to provide additional lighting beyond the ambient conditions, ensuring optimal visibility during printing. To control the printer’s spatial movements, extrusion flow rate, lighting, camera operation, and data collection or processing, a Raspberry Pi Model 4B was selected as the microprocessor. The processor orchestrates the printer’s operations through a master-slave communication setup, where commands are transmitted from the microprocessor to the printer and other components. In terms of power supply requirements, the printer operates on a main power supply of 24 V DC, while a separate 5 V DC 3A power supply is used to power the Raspberry Pi microprocessor, ensuring reliable operation and coordination of the printer’s functions.

### Bioprinting workflow and quality metrics

To monitor the quality of bioink extrusion we developed a print path that begins with an undistributed horizontal straight line, followed by a vertical direction change at a specific distance from the initial line (Fig 1c). Subsequently, three horizontal straight lines, each progressively closer to the initial undistributed line, are printed to observe co-printing disruption at varying proximities. Additionally, for illustrative purposes, the path continues at an increased speed without extrusion while moving in parallel to the initial line. It then vertically diverges before returning to intersect the initial line. Finally, it returns to horizontal movement and completes the loop. The print path was created using computer-aided design (CAD) software in the form of a cuboid filament with a width and height of 285 microns. The STL file was subsequently processed by a slicing software, converting it into the machine-readable G-code language. The G-code was then sent line-by-line to the pre-existing printer controller through the microcontroller while executing a Python script. All prints were performed at an ambient temperature of 25°C, a printing speed of 1.5 mm/s, and a travelling speed of 6.5 mm/s, using a straight dispensing tip of 25 gauge with a nominal inner diameter of 254 microns. Prints were performed at a z-height of 1.1mm from the bottom of the glass dish housing the support hydrogel. A brightfield microscope was used to accurately measure the outer diameter of the needle of the nozzle tip with the average value being 428 microns.

### Statistical analysis

GraphPad Prism 10 software was used for all statistical analyses, and Microsoft Excel was used for data handling. Statistical comparisons between experimental groups were performed using a two-way ANOVA with Tukey post hoc testing. Sample distribution was assumed normal with equal variance.

